# ER bodies are induced by *Pseudomonas syringae* and negatively regulate immunity

**DOI:** 10.1101/2020.11.22.393215

**Authors:** José S. Rufián, James M. Elmore, Eduardo R. Bejarano, Carmen R. Beuzon, Gitta L. Coaker

## Abstract

ER bodies are endoplasmic reticulum-derived organelles present in plants belonging to the *Brassicales* order. In *Arabidopsis thaliana*, ER bodies are ubiquitous in cotyledons and roots, and present only in certain cell types in rosette leaves. However, both wounding and jasmonic acid treatment induce the formation of ER bodies in leaves. Formation of this structure is dependent on the transcription factor *NAI1*. The main components of the ER bodies are β-glucosidases (BGLUs), enzymes that hydrolyze specialized compounds. In *Arabidopsis*, PYK10 (BGLU23) and BGLU18 are the most abundant ER body proteins. In this work, we found that ER bodies are downregulated as a consequence of the immune responses induced by bacterial flagellin perception. *Arabidopsis* mutants defective in ER body formation show enhanced responses upon flagellin perception and enhanced resistance to bacterial infections. Furthermore, the bacterial toxin coronatine induces the formation of *de novo* ER bodies in leaves and its virulence function is partially dependent on this structure. Finally, we show that performance of the polyphagous beet armyworm herbivore, *Spodoptera exigua*, increases in plants lacking ER bodies. Altogether, we provide new evidence for the role of the ER bodies in plant immune responses.

## INTRODUCTION

ER bodies are rod-shaped endoplasmic reticulum (ER)-associated structures that are present in three families within the Brassicales order (Nakano et al., 2014a). ER bodies are constitutively present in roots and cotyledons of young plants (constitutive ER bodies), but their abundance decreases with senescence (Matsushima et al., 2002). In *Arabidopsis* rosette leaves, ER bodies are constitutively present in marginal cells, epidermal cells covering the midrib and extremely large pavement cells (Leaf ER bodies) (Nakazaki et al., 2019b). Furthermore, the formation of ER bodies in rosette leaves is induced by Methyl Jasmonate (MeJA) treatment or wounding (Inducible ER bodies) (Matsushima et al., 2002).

Formation of both constitutive and leaf ER bodies is dependent on the basic-helix-loop-helix transcription factor *NAI1* (Matsushima et al., 2004) and the essential protein *NAI2* (Yamada et al., 2008). Inducible ER bodies are formed in a *nai1* mutant, but differin shape and composition with the ER bodies induced in wild-type Col-0 (Ogasawara et al., 2009). The main components of the constitutive ER bodies are the β-glucosidases (BGLU) PYK10 (BGLU23), BGLU21 and BGLU22 (Matsushima et al., 2003), while inducible-ER bodies also contain BGLU18 (Ogasawara et al., 2009). These myrosinases belong to a subfamily of 8 β-glucosidases (*i. e*. BGLU18 to BGLU25) (Xu et al., 2004) with ER-retention signals (Yamada et al., 2011). Leaf ER bodies mainly contain PYK10 and BGLU18 (Nakazaki et al., 2019b). PYK10 has been described as an atypical myrosinase that hydrolyzes Indole Glucosinolates (IG) (Nakano et al., 2017). Upon tissue damage, PYK10 associates in the cytosol with PYK10-binding protein 1 (PBP1), activating PYK10 function (Nagano et al., 2005). An *Arabidopsis pyk10/blgu18* double mutant accumulates high amounts of the IG 4-methoxyindol-3-ylmethylglucosinolate (4MI3G) after tissue damage (Nakazaki et al., 2019b). IGs and the products resulting from their breakdown have a role in the defense responses against microbial pathogens and herbivory (Bednarek et al., 2009; Clay et al., 2009; Millet et al., 2010; Müller et al., 2010; Johansson et al., 2014; Nakazaki et al., 2019b). IGs are also required for the establishment of beneficial interactions with the endophytic fungus *Piriformospora indica* by appropriately regulating the level of root colonization through *PYK10* (Sherameti et al., 2008; Lahrmann et al., 2015). Collectively, multiple independent studies have supported a role in defense responses for ER bodies (Yamada et al., 2011; Nakano et al., 2014b).

Plants are constantly exposed to a variety of biotic and abiotic stresses. Surface localized plant pattern-recognition receptors (PRRs) act as one of the first lines of defense against pathogens and are able to recognize conserved pathogen features, such as flagellin, as non-self (Couto and Zipfel, 2016). One of the best-characterized PRRs is the flagellin receptor FLS2 (Flagellin Sensing 2) which recognizes flg22, a 22 amino acid epitope of bacterial flagellin (Gómez-Gómez and Boller, 2000). Perception of flg22 by FLS2 induces a number of cellular events resulting in Pattern-Triggered Immunity (PTI), which restricts pathogen proliferation. Some of the cellular events that take place during PTI are the activation of MAP Kinases (MAPK) and calcium-dependent protein kinases, the production of reactive oxygen species (ROS), stomatal closure and the formation of callose (β-1,3-glucan) deposits in the cell wall (Henry et al., 2013). However, successful pathogens have evolved different mechanisms to suppress defense responses and successfully colonize their host.

*Pseudomonas syringae* is a Gram-negative bacterium that has been extensively used to study plant immune signaling. *P. syringae* virulence depends on the Type III Secretion System (T3SS), a complex nanomachine that translocates proteins, called effectors, into the cytosol of the plant cell. Most of the effectors suppress PTI acting on different cellular pathways (Macho, 2015). Another virulence mechanisms used by bacterial pathogens to subvert plant immunity is the production of phytotoxins. *P. syringae* pv. tomato DC3000 (*Pst* DC3000) produces coronatine, a polyketide toxin that acts as a molecular mimic of the plant hormone jasmonic acid (JA) (Weiler et al., 1994). Coronatine induces stomata re-opening upon PRR perception to allow bacterial colonization (Melotto et al., 2006; Melotto et al., 2008; Gimenez-Ibanez et al., 2017), and also suppresses callose deposition and induces bacterial growth within the apoplast (Geng et al., 2012).

In this work, we investigated the biological function of ER bodies during plant immunity. We found that ER bodies are downregulated after flg22-triggered immune activation but induced after *P. syringae* infection in a coronatine-dependent manner. Furthermore, loci underlying ER body formation act as negative regulators of immunity against bacterial pathogens but are required for a complete response against the chewing insect *Spodoptera exigua*. With these results, we report a new role of ER bodies in virulence of phytopathogenic bacteria and their suppression by plant immune perception..

## RESULTS

### ER bodies are rapidly inhibited upon flagellin perception

To investigate the role of ER body formation after plant pathogen perception, we used an *Arabidopsis* transgenic line expressing GFP with the ER-retention signal HDEL (GFP-HDEL), which allows the visualization of ER bodies (Mitsuhashi et al., 2000). We vacuum-infiltrated 2 week-old *Arabidopsis* plants with either 10 μM flg22 or water and visualized cotyledons three hours later by confocal microscopy. We were able to detect and quantify the ER bodies in cotyledons, where this structure is constitutively present, in mock-treated plants. However, after flg22 treatment, only a small number of ER bodies were detected (Fig. 1A). This decrease in ER body detection was statistically different compared to water-treated controls (Fig. 1A). To further investigate the disappearance of ER bodies within the same tissue, we monitored cotyledons for 120 minutes after either water (mock) or flg22 treatment. ER bodies lose their characteristic rod-shape within 60 minutes post-flg22 treatment, and the GFP signal is diffused throughout the cell at later time points (Fig. 1B).

**Figure 1.**
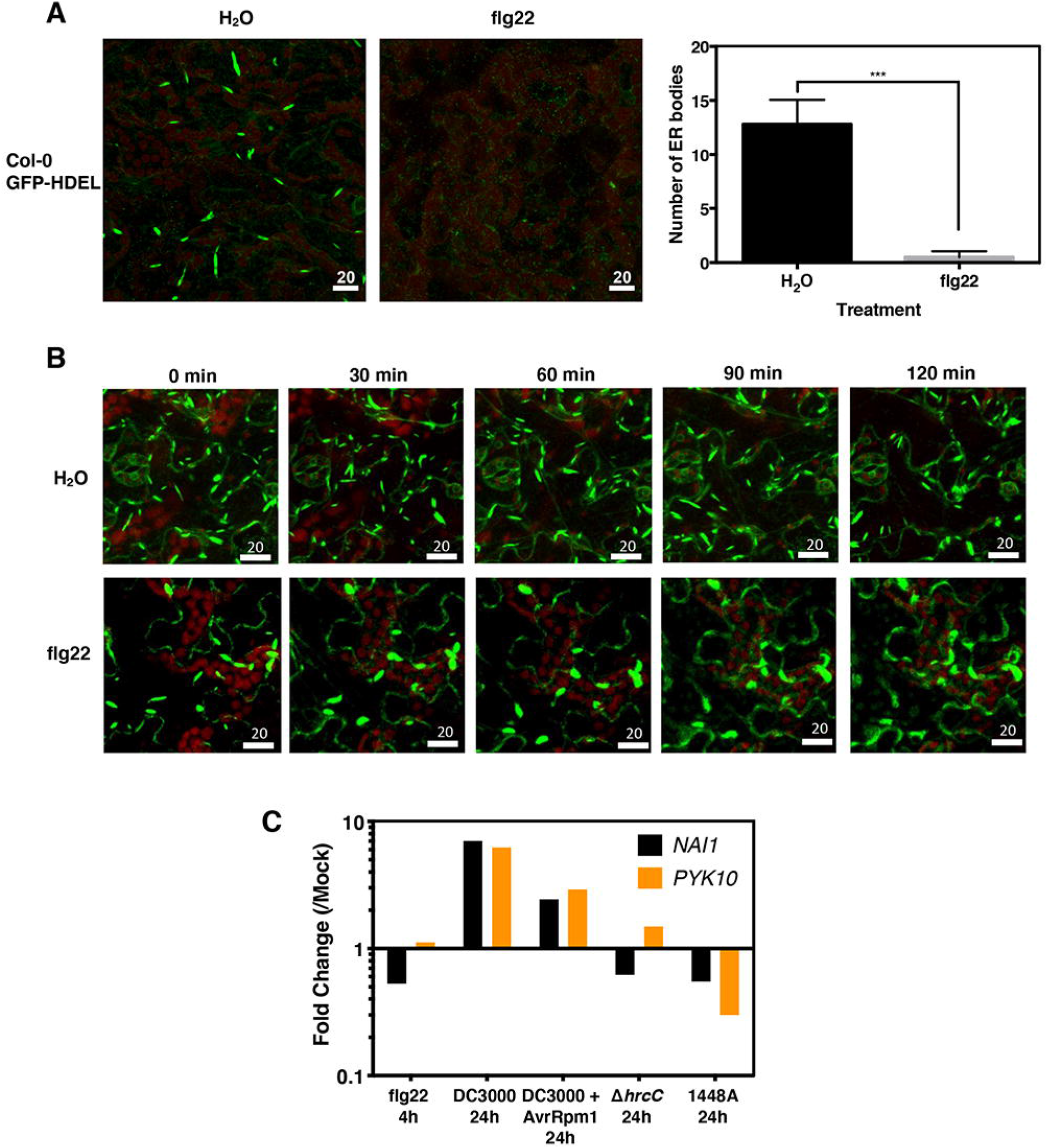
ER bodies are downregulated after activation of PTI. (**A**) Constitutive ER bodies disappear after flg22 treatment. Two-week-old *Arabidopsis* GFP-HDEL plants were vacuum infiltrated with water or 100 nM flg22. Cotyledons were observed in a Zeiss LSM710 confocal microscope three hours after treatment. Red autofluorescence correspond to chloroplasts. For ER body quantification, three fields from three independent cotyledons were used. ER bodies were quantified using Fiji. The experiment was repeated four times with similar results and a mean value ± SE of all quantifications is shown. Statistical differences were determined with a two-tailed t-test comparing water treatment with flg22. Three asterisks indicate p<0.001. Scale bar: 20 µm. (**B**) Two-week-old *Arabidopsis* GFP-HDEL plants were vacuum infiltrated with water or 100 nM flg22. The infiltrated cotyledons were monitored for 120 minutes taken photos every 5 minutes with a Leica SP8 confocal microscope. (**C**) Expression patterns of *nai1* and *pyk10* genes under different biotic stress. Data was obtained from available microarray in eFP Browser. Five week-old Col-0 plant leaves were syringe-infiltrated with either a 1 μM flg22 solution (using water as mock treatment) or the indicated bacterial strain at 10^8^ cfu/ml (using 10 mM MgCl_2_ as mock treatment). Samples were taken 24 hours after treatment. Fold change relative to mock treatment is shown.

To analyze the impact of the flg22 treatment in ER body formation at the transcriptional level, we analyzed the transcriptional expression of *NAI1* and *PYK10* under different biotic treatments using an available dataset (NASCArrays-120). We found that flg22 treatment reduces the expression level of *NAI1* and *PYK10* in an *FLS2*-dependent manner (Fig. 1C, Fig. S1), pointing to regulation of ER bodies at the transcriptional level as a consequence of activation of defense responses. However, inoculation with the virulent bacterial strain *Pst* DC3000 induces the expression of both *NAI1* and *PYK10* at 24h, and the same occurs with a strain expressing the avirulence effector AvrRpm1 (Fig. 1C). Interestingly, both the T3SS DC3000 mutant derivative Δ*hrcC* and the non-host strain 1448A reduced the accumulation of *NAI1* transcripts (Fig. 1C). Altogether, these results show a downregulation of ER bodies as a consequence of the activation of flg22-triggered immunity and suggest a role of this subcellular structure during the defense response against *P. syringae*.

### *Arabidopsis* mutants impacting ER body formation and contents exhibit enhanced immune responses against bacterial pathogens

After flg22 perception, many cellular events take place to restrict pathogen invasion, including activation of MAPKs and calcium dependent protein kinases minutes after perception, followed by ROS production, activation of defense gene expression and several hours later, callose deposition (Henry et al., 2013). To further characterize the role of the ER bodies in flg22-induced responses, we measured ROS production as an early event of PTI, in plants affected in the formation of ER bodies. We treated 3-week-old Col-0, *nai1-1, pyk10-1* and *pyk10-1/blgu21* plants, with 100 nM flg22 and measured the ROS burst. The total ROS produced in all the mutant backgrounds tested was higher than the ROS produced in Col-0 (Fig. 2A). This increase in ROS production in response to flg22 also occurs in both *bglu18-1* and *pbp1-1* mutants (Fig S2A). To analyze a late event of PTI, we treated four-week-old plants with flg22 and quantified callose deposition. Consistent with the ROS burst, the number of callose deposits was two times higher in ER bodies mutants compared to Col-0 (Fig 2B). These data indicate that a loss of ER bodies or its main components induce an enhanced response to bacterial flg22, pointing to a negative role of ER bodies on PTI.

**Figure 2.**
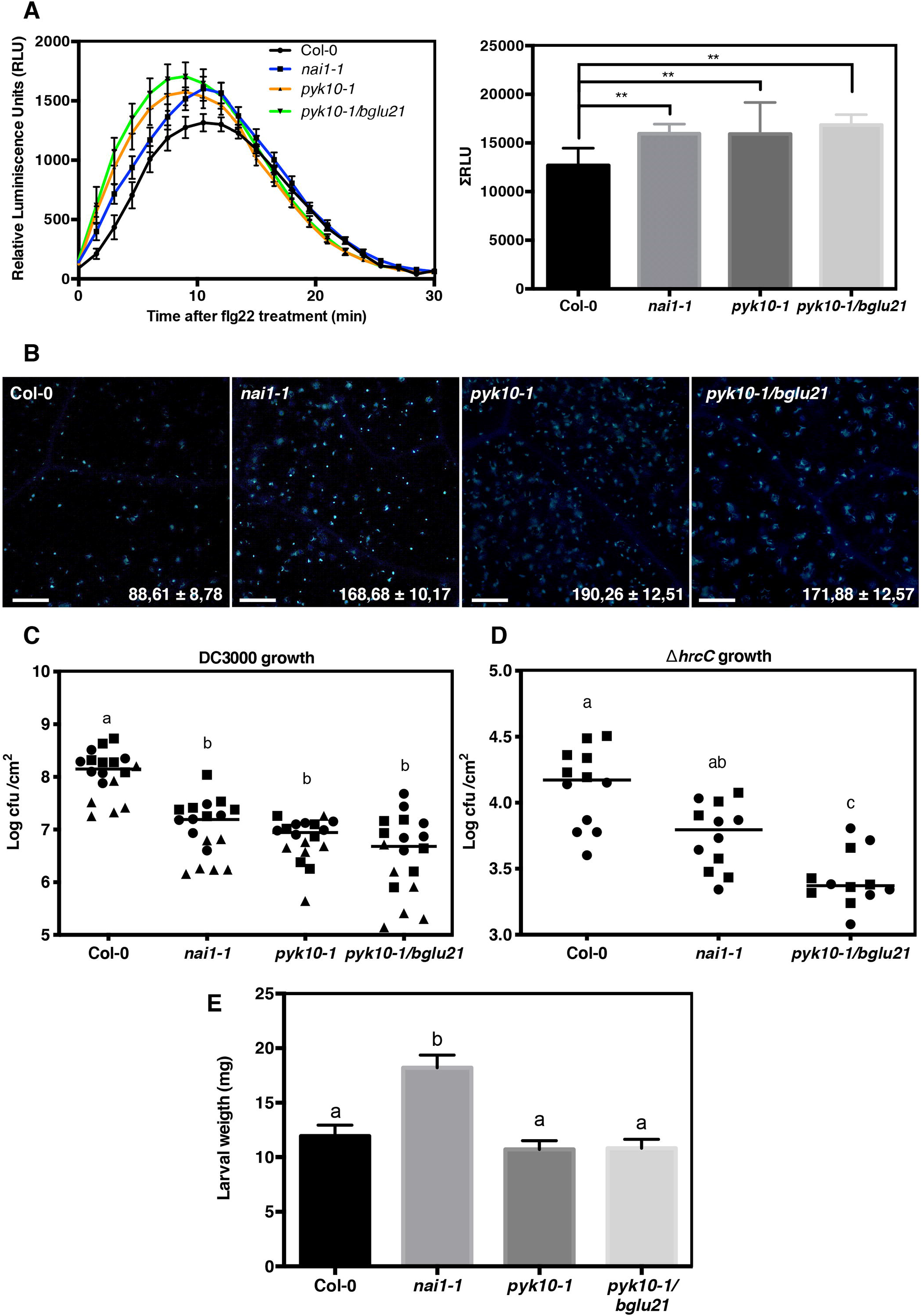
ER body mutants exhibit enhanced resistance against *P. syringae*. (**A**) flg22-induced ROS burst in Col-0, *nai1-1, pyk10-1* and *pyk10-1/bglu21*. Three-week-old *Arabidopsis* leaf discs were treated with 100 nM flg22 and ROS was quantified using a luminescence based assay. The left graph represent the dynamics of the ROS produced in the different genotypes and the right graph represent the total relative lights units (RLU) detected over a 30 minutes period. Error bars indicate SD, *n* = 16. Statistically differences were determined with one-way ANOVA (α = 0.05) with Tukey’s multiple comparisons test and different letters indicate statistical significance. The experiment was repeated four times with similar results and a representative experiment is shown. (**B**) Quantification of flg22-induced callose deposition in Col-0, *nai1-1, pyk10-1* and *pyk10-1/bglu21*. Four-week-old plants were infiltrated with 10 µM flg22 and callose deposits were visualized under an epifluorescence microscope eighteen hours after treatment. Four images were taken per leaf and nine leaves were used per experiment. The mean number of deposits was quantified and is included ± SE in the bottom-right corner of each representative image. The experiment was repeated three times with similar results, and a representative experiment is shown. White bar: 100 µm. (**C**) Growth of *P. syringae* pv. *tomato* DC3000 in Col-0, *nai1-1, pyk10-1* and *pyk10-1/bglu21*. Five-week-old plant leaves were syringe inoculated with a suspension of 5×10^4^ CFU/mL. Four days after inoculation, bacteria were recovered and quantified. Values from three independent replicates are shown (*n*=18). Different symbols represent individual values from different replicates. Statistically differences were determined with by ANOVA (α = 0.05) with Tukey’s multiple comparisons test and different letters indicate statistical significance. (**D**) Growth of the *P. syringae* pv. *tomato* Δ*hrcC* derivative in Col-0, *nai1-1*, and *pyk10-1/bglu21*. Five-week-old plant leaves were syringe inoculated with a suspension of 5×10^4^ CFU/mL. Four days after inoculation, bacteria were recovered and quantified. Values from three independent replicates are shown (*n*=18). Different symbols represent individual values from different replicates. Statistically differences were determined by ANOVA (α= 0.05) with Tukey’s multiple comparisons test and different letters indicate statistical significance. (E) Three-week-old *Arabidopsis* Col-0, *nai1-1, pyk10-1* and *pyk10-1/bglu21* plants were challenged with three-days-old *S. exigua* larvae. Seven days after feeding, fresh weight of each larva was measured. Bars represent the mean weight ± SE (*n*=20). Statistically differences were determined with one-way ANOVA (α= 0.05) with Tukey’s multiple comparisons test and different letters indicate statistical significance. The experiment was repeated three times with similar results, and a representative experiment is shown.

To determine the role of the ER bodies in bacterial growth, we quantified *Pst* DC3000 titers in Col-0 and in the different mutant backgrounds (Fig. 2C). *Pst* DC3000 exhibited decreased growth (>1 log) in the *nai1-1* mutant compared to Col-0. Interestingly, both the *pyk10-1* and the *pyk10-1/bglu21* mutants exhibited a stronger decrease in bacterial growth, with 1.5 Log reduction compared to wild-type Col-0 (Fig 2C). No differences were detected between *pyk10* and *pyk10-1/bglu21*, indicating that *PYK10* is a critical gene involved in negative regulation of bacterial growth. Plants lacking *PBP1*, necessary for PYK10 function, or *BGLU18*, the main component of inducible ER bodies, also showed enhanced resistance to bacterial infection (Fig, S2B).

A critical component controlling bacterial virulence is the ability to deliver effectors through the type III secretion system (Macho, 2015). In order to investigate the relationship between immune suppression and ER body formation, we used DC3000 Δ*hrcC*, which is unable to translocate effectors to the plant cell, and thus, does not suppress PTI. We measured the growth of Δ*hrcC* in ER body mutants. In the *nai1-1* background, the growth of Δ*hrcC* was not significantly different from that in Col-0. However, we observed a slight but significant decrease in growth in the *pyk10-1/bglu21* background (Fig. 2D). The increased *P. syringae* growth restriction in *pyk10-1* and *pyk10-1/bglu21* backgrounds compared with *nai1* (Fig. 2C and 2D) could indicate a NAI1-independent regulation for *PYK10* and/or *BGLU21*. Altogether, these results show that the absence of ER bodies increases resistance against the bacterial pathogen *P. syringae*.

### *NAI1* contributes to the defense response against *S. exigua*

ER bodies have been proposed to participate in the defense response against herbivores (Yamada et al., 2011; Nakano et al., 2014b). A recent study demonstrated increased susceptibility of the double mutant *bglu18 pyk10* to the terrestrial isopod *Armadillidium vulgare* (Nakazaki et al., 2019b). The *pyk10/bglu21* mutant forms ER bodies, although they are larger and less numerous than those in wild-type plants (Nagano et al. 2009). We measured the performance of the generalist leaf-chewing herbivore *S. exigua* in different ER body related mutants. Performance assays have been widely used to study the defense mechanisms against herbivory (Cipollini et al., 2004; Chung et al., 2008; Van Oosten et al., 2008; Müller et al., 2010; Santamaría et al., 2017). We fed *S. exigua* larvae with Col-0 and the ER body mutants *nai1-1, pyk10-1* and *pyk10-1/bglu21* plants, and measured the weight of the larvae after 7 days. The fresh weight of *S. exigua* larvae was higher in *nai1-1* plants compared with the Col-0 control, confirming a role of ER bodies in the defense response against herbivore insects. Surprisingly, the performance of *S. exigua* in either *pyk10-1* or *pyk10-1/bglu21* plants was not different compared Col-0, suggesting a redundant role of *PYK10* and *BGLU21* with other myrosinases in this context. (Fig. 2E)

### Coronatine induces the formation of ER bodies in rosette leaves

Transcriptomic analyses indicate that inoculation with virulent *Pst* DC3000 induces the expression of *NAI1* and *PYK10* (Fig. 1C). To check if this increased expression results in *de novo* formation of ER bodies, we inoculated rosette leaves of *Arabidopsis* GFP-HDEL plants with a relatively high inoculum (5×10^7^ CFU/mL) of *Pst* DC3000, the derivative Δ*hrcC* and a coronatine mutant (*cor*-, *Pst* DC3118). We applied the bacterial suspension using a brush to avoid wounding tissue. Three days after inoculation, we observed the samples under the confocal microscope (Fig. 3A). No ER bodies were found in the mock-treated leaves, in agreement with the absence of ER bodies in certain rosette leaves (Matsushima et al., 2002; Nakazaki et al., 2019b). Interestingly, a large number of ER bodies was found in leaves inoculated with wild-type *Pst* DC3000. The distribution of the ER bodies was patchy, suggesting that only some cells were forming the structure. To investigate if the formation of ER bodies localized surrounding bacterial colonies, we generated a *Pst* DC3000 derivative tagged with eYFP. Three days after inoculation, we observed a large number of ER bodies surrounding the bacterial microcolony in Col-0 (Fig. 3B,C) but only a few aberrant ER bodies in the *nai1-1* mutant (Fig. 3C), suggesting that ER bodies induced by *Pst* DC3000 are dependent on *NAI1*. To determine which bacterial factors are inducing the formation of ER bodies in rosette leaves, we inoculated Col-0 GFP-HDEL leaves with the Δ*hrcC* derivative and the *cor*-mutant, both tagged with eYFP. We were unable to detect Δ*hrcC* – eYFP microcolonies due to their extremely small size, consistent with their reduced growth *in planta*. Nevertheless, we did not detect ER bodies in leaves inoculated with the Δ*hrcC* mutant (Fig. 3A). Interestingly, the *cor-*eYFP mutant was unable to induce the formation of *de novo* ER bodies, consistent with the role of coronatine mimicking JA (Fig. 3). This result indicates that induction of ER bodies could be a virulence function of coronatine. Analyzing transcriptomic data of plants inoculated either with *Pst* DC3000 or the *cor*-mutant, we observed that many genes related to ER bodies, including *NAI1, PYK10* and *BGLU18*, were upregulated by *Pst* DC3000 but not after inoculation with *Pst* DC3118 (*cor*-) (Fig. S3). To investigate the ability of coronatine to promote bacterial virulence in the absence of wild-type ER body formation, we measured the growth of *Pst* DC3000 and the *Pst cor-*mutant in Col-0, *nai1-1, pyk10-1* and *pyk10-1/bglu21*. In order to bypass the role of coronatine on re-opening stomata (Melotto et al., 2006; Melotto et al., 2008), we performed syringe inoculation in five-week old plant leaves. Four days after inoculation, a strong growth restriction was observed in the coronatine mutant compared with wild-type *Pst* DC3000 in Col-0 (Fig. 3D). In the ER body-defective backgrounds, a stronger growth restriction was observed for *Pst* DC3000 (Fig. 2C, Fig. 3D). However, the growth of the coronatine mutant was not significantly different between wild-type Col-0 and the ER body mutants.

**Figure 3.**
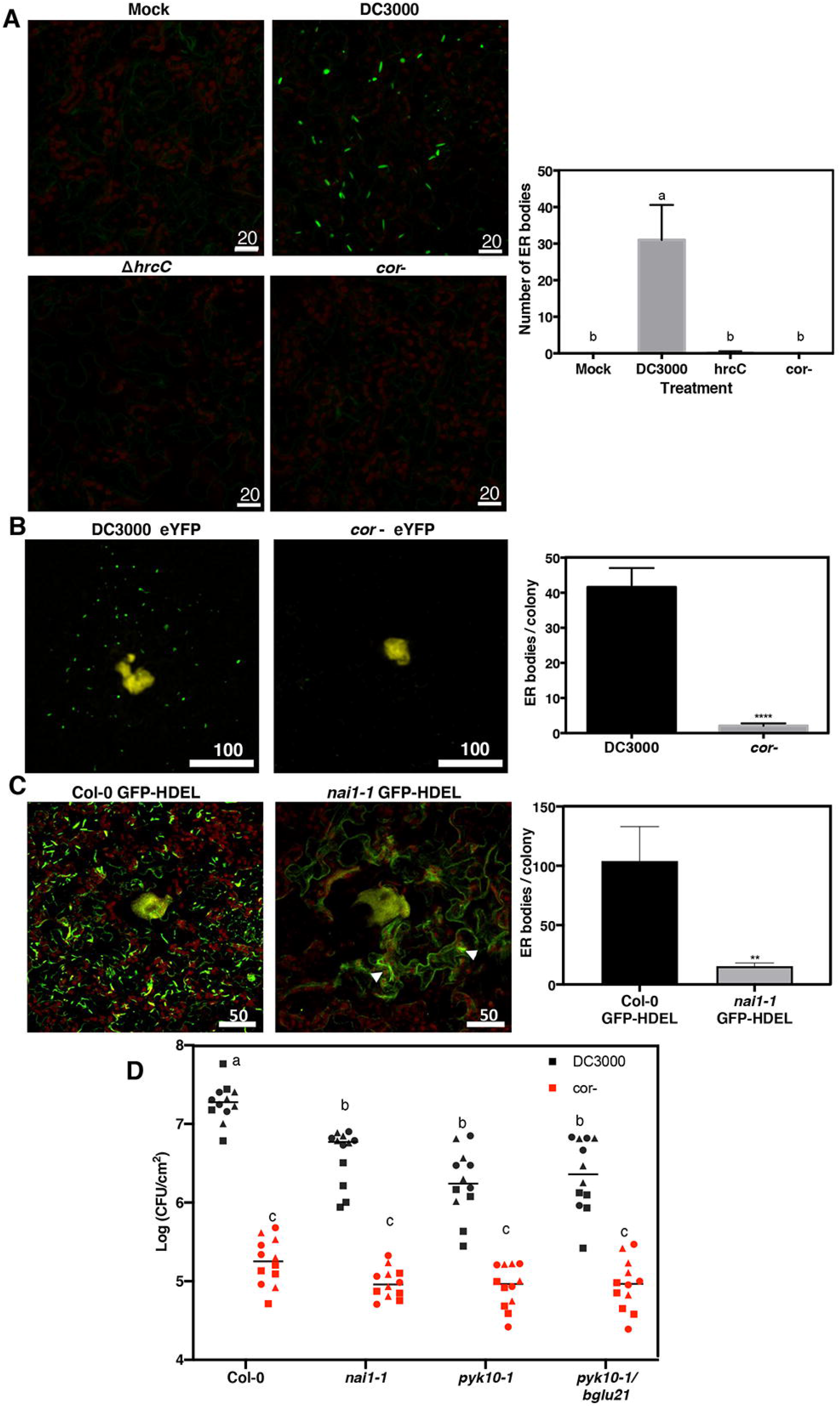
Coronatine induces formation of ER bodies in leaves and its virulence function is compromised in mutants impaired in ER body formation. (**A-C**) Confocal microscope images of bacterial micro-colonies (yellow) and ER bodies (green). Five-week-old *Arabidopsis* GFP-HDEL plants were brush-inoculated with a suspension containing 5×10^7^ CFU/mL (A) or syringe inoculated with a bacterial suspension containing 5×10^4^ CFU/mL (B, C). Three days after inoculation, leaf sections were visualized under the confocal microscope. A representative z-stack projection image is shown. ER bodies were quantified from the green channel using Fiji. Bars represent the mean number of ER bodies in (A) or the number of ER bodies surrounding a single bacterial micro colony in (B, C) ± SE. Red autofluorescence correspond to chloroplasts. The experiments were repeated three times with similar results, and the mean of all experiments is shown. ER bodies were quantified using Fiji and statistically differences were determined with one-way ANOVA (α= 0.05) with Tukey’s multiple comparison test in (A) and different letters indicate statistical significance. In (B, C) data were analyzed by a t-test (α= 0.001). White bar: 20 µm (A), 100 µm (B), 50 µm (C). (**D**) Growth of *P. syringae* pv. *tomato* DC3000 and its derivative coronatine mutant (*cor* -, DC3118) in Col-0, *nai1-1, pyk10-1* and *pyk10-1/bglu21*. Five-week-old plant leaves were syringe inoculated with a bacterial suspension of 5×10^4^ CFU/mL. Four days after inoculation, bacteria were recovered and quantified. Values from three independent replicates are shown (*n*=12). Different symbols represent individual values from different replicates. Statistically differences were determined by ANOVA (α = 0.05) with Tukey’s multiple comparisons test and different letters indicate statistical significance.

## DISCUSSION

The main contents in ER bodies are β -glucosidases, enzymes which can produce toxic compounds to protect the plant from herbivore attack (Matsushima et al., 2003). Induction of ER bodies by wounding or by MeJA treatment suggests an association with plant defense (Nakano et al., 2014b). However, there are few experimental evidences of the association between ER bodies and plant immunity. Plants are infected by diverse pathogens that often utilize opposing virulence strategies and are differentially impacted by plant immune responses (e.g. chewing insects versus biotrophic/hemibiotrophic pathogens). In this study we demonstrate contrasting roles of ER bodies in the plant response to pathogenic bacteria and herbivores.

Previous studies have implicated ER bodies in plant protection against fungi and feeding insects. Sherameti et al. (2008) demonstrated that *PYK10* is required to protect plants against excessive root colonization by the endophytic fungus *Piriformospora indica*. Furthermore, Nakazaki et al. (2019b) showed that *bglu18 pyk10* double mutant plants are more susceptible than wild-type to the attack of the terrestrial isopod *Armadillidium vulgare* (pill-bug). In this work, we experimentally link the function of the ER bodies with the plant response against the bacterial pathogen *P. syringae* and the herbivore *S. exigua*. We found an increased weight acquisition of the generalist chewing herbivore *S. exigua* in *nai1-1* plants compared with Col-0, but not in *pyk10-1* and *pyk10-1/bglu21* mutants (Fig. 2E). *S. exigua* feeds on leaves, where constitutive ER bodies are absent (Matsushima et al., 2002). Constitutive leaf ER bodies, which are dependent on *NAI1*, localize in the edges of the leaves and mainly contain PYK10 and BGLU18 (Nakazaki et al., 2019b). The biogenesis and content of leaf ER bodies is regulated by both Jasmonic Acid (JA) dependent and independent pathways (Nakazaki et al., 2019a). While *NAI1* expression, and therefore PYK10 accumulation is dependent on JA, BGLU18 accumulation in leaf ER bodies is independent on JA (Nakazaki et al., 2019a). Since the performance of *S. exigua* is not affected in *pyk10-1* mutant, the genetic analysis points to *BGLU18* as the responsible of the defense response against *S. exigua*. Furthermore, chewing induces JA accumulation (Rehrig et al., 2014) and this signal would potentially lead to the formation of the inducible ER bodies. The *nai1* mutant is compromised in formation of inducible local and systemic ER bodies. In the *nai1* mutant, inducible ER bodies are long and tubular and mainly observed in cells surrounding the wound site (Ogasawara et al., 2009). We also observed that *Pst* DC3000 was unable to induce ER body formation in the *nai1* mutant, indicating that the absence or altered composition/structure of ER bodies in *NAI1* suppresses growth of the hemibiotrophic pathogen *P. syringae*, while enhancing performance of a chewing herbivore.

*Pst* DC3000 induces the formation of ER bodies in leaves. Since the flagellin peptide flg22 downregulates the formation of constitutive ER bodies and we have shown that ER bodies negatively regulate PTI, we hypothesize that *P. syringae*-induction of ER bodies is a virulence strategy to suppress host immune responses. The two main virulence factors of *P. syringae* pv. tomato are the phytotoxin coronatine and the T3SS effectors. The bacterial phytotoxin coronatine, which mimics the function of JA in the plant (Bender et al., 1999), induces the formation of ER bodies in leaves (Fig 3A,B). JA is perceived in the plant through degradation of JASMONATE ZIM-DOMAIN (JAZ) transcriptional repressors and JAZ proteins suppress the expression of ER body components as well as indole glucosinolates (Guo et al., 2018; Ghorbel et al., 2021). Thus, JA or pathogen JA-mimics can regulate the abundance of this specialized organelle to induce or suppress defense depending on pathogen type. The most studied coronatine virulence function is to re-open stomata upon PAMP-induced closure (Melotto et al., 2006; Melotto et al., 2008). However, coronatine also has been shown to suppress immunity once the bacteria colonize the apoplast (Fig. 3C; (Geng et al., 2012)). The T3SS defective mutant Δ*hrcC* is not able to induce the formation of ER bodies, which would indicate an effect of T3Es on the induction of ER bodies. However, we cannot rule out that the absence of ER bodies is due to an insufficient amount of coronatine produced by the Δ*hrcC* microcolonies.

Bacterial flg22 perception leads to a down regulation of the ER bodies at both transcriptional and post-transcriptional levels (Fig. 1 and S1). We found that *Arabidopsis* mutants defective in either ER body biogenesis or composition are more resistant to pathogenic bacteria (Fig. 2 and S2). The main components of ER bodies are myrosinases, enzymes that modify glucosinolates to initiate the glucosinolate breakdown, which will result in different products depending on the biochemical properties of the modifier protein and the chemical nature of the glucosinolate side chain (Wittstock and Burow, 2010; Sugiyama and Hirai, 2019). Those products can function as defense compounds or as signaling molecules.

PYK10/BGLU23 is an atypical myrosinase similar to PENETRATION 2 (PEN2, BGLU26) that hydrolyzes indole glucosinolates (IGs) (Nakano et al., 2017; Nakazaki et al., 2019b). PEN2-mediated conversion of 4-methoxyindol-3-ylmethylglucosinolate (4MI3G) into glucosinolate-derived products plays an important role in the defense response against fungi (Bednarek et al., 2009). Interestingly, *PEN2* is required for callose deposition upon flg22 perception and for resistance against *Pst* DC3000 infection in seedlings grown in liquid media (Clay et al., 2009; Johansson et al., 2014). In contrast, in adult plants grown in soil, callose deposition and *Pst* DC3000 growth is not affected in the *pen2-1* mutant (Geng et al., 2012). The *pen2-1* mutant grown in soil exhibits enhanced resistance to *P. syringae* pv. *maculicola* (Stahl et al., 2016). These apparently contradictory results indicate that PEN2 activity is dependent on age and/or the metabolic state of the plant. 4MI3G overaccumulates in homogenized tissue of *bglu18/pyk10* mutants indicating that one or both of these β-glucosidases hydrolyze 4MI3G (Nakazaki et al., 2019). Our results indicate that the single *pyk10-1* mutant and the double *pyk10-1/bglu21* mutant accumulate an increased number of callose deposits in response to flg22 treatment and are more resistant to *Pst* DC3000. Although both PEN2 and PYK10 seem to have the same substrate (4MI3G), their respective mutants differ on phenotypes such as flg22-induced callose deposition and resistance to *P. syringae Pst* DC3000 (Clay et al., 2009; Johansson et al., 2014). Apart from the above-mentioned effect related to growth conditions, one of the reasons explaining these differences could be the different compartmentalization of both PEN2 and PYK10. While PEN2 is localized in peroxisome and mitochondria (Lipka et al., 2005; Fuchs et al., 2016), PYK10 localizes in ER bodies (Matsushima et al., 2003). Furthermore, myrosinase activity is regulated by different Myrosinase-Biding Proteins and Myrosinase-Associated Proteins, as well as different specifier proteins, which lead to the formation of different breakdown products (Wittstock et al., 2016). This variety of compounds can have a direct or indirect effect on the plant defense response (Wittstock et al., 2016). It is possible that mutants lacking ER bodies or core components of ER bodies also exhibit heightened basal defense, resulting in enhanced ROS and callose production.

In conclusion, we have demonstrated the involvement of ER body-dependent regulation of immunity to bacteria and induction of ER bodies as a pathogen virulence strategy. Further work will shed light on how this novel function of ER bodies influences bacterial-plant interactions and pave the way for the discovery of the role of indole-glucosinolates breakdown on the response against bacterial invasion in plants.

### Experimental procedures

#### Plant material and growth conditions

*A. thaliana* Col-0, the transgenic line expressing GFP-HDEL (Hayashi et al., 2001) and the mutants *nai1-1* (Matsushima et al., 2003), *pyk10-1* (Nagano et al., 2008), *pyk10-1/bglu21* (Nagano et al., 2009), *bglu18-1* (Ogasawara et al., 2009) and *pbp1-1* (Nagano et al., 2005) were grown in a controlled environment chamber at 23°C, 70% relative humidity, and a 10h/14h light/dark photoperiod with light intensity of 100µE.m^-2.^S^-1^. All seeds were stratified for 3-4 days at 4°C before sowing into soil.

#### Bacterial strains, growth conditions and pathogen assays

*Pseudomonas syringae* pv. *tomato* DC3000 (Cuppels, 1986) and the derivative strains Δ*hrcC* and *cor* ^*-*^ (cma-cfa-; (Brooks et al., 2004)) were grown at 28°C in Lysogeny Broth (LB) medium (Bertani, 1951). Antibiotics were used when appropriate, at the following concentrations: Rifampicin (100 μg/ml), kanamycin (25 μg/ml) and gentamycin (10 μg/ml).

The *Pst* DC3000, Δ*hrcC* and *cor* ^*-*^ eYFP (enhanced yellow fluorescent protein) derivatives were generated using a Tn7 delivery system as previously described (Rufian et al., 2018)

For pathogen assays, five-week-old *Arabidopsis* plants were inoculated with a *P. syringae* suspension of 5×10^4^ CFU/mL prepared in 10 mM MgCl_2_. Three leaves per plant were syringe infiltrated and five plants were used for each experiment. Four days after inoculation, three leaf discs of 1 cm of diameter (one per leaf were ground in 1 mL 10 mM MgCl_2,_ and serial dilutions were plated.

#### Transcriptomic analyses

For the analysis of NAI1 and PYK10 expression under different biotic stresses, we used transcriptomic data available in eFP Browser. Specifically, we used the dataset AtGenExpress: Response to virulent, avirulent, type III□secretion system deficient and nonhost bacteria (NASCArrays-120). 5-week-old Col-0 plants were infiltrated with the corresponding *P. syringae* strain. Samples were taken 24h after treatment and RNA was isolated and hybridized to the ATH1 GeneChip. The data were normalized by GCOS normalization, TGT 100.

#### flg22-related assays

For ROS burst assays, leaf discs were collected using a cork borer of 5mm diameter from three-week-old *Arabidopsis* plants and floated overnight in deionized water. The next day, the water was replaced with an assay solution containing 17 mg/ml luminol (Sigma), 10 mg/ml horse radish peroxidase (Sigma) and 100 nM flg22 (Genscript). Luminescence was measured using Tristar multimode reader (Berthold technology).

Callose deposits were quantified from four-week-old *Arabidopsis* leaves infiltrated with 10 μM flg22 solution. Eighteen hours after treatment, leaves were detached and incubated 15 minutes at 65°C in 5 ml of alcoholic lactophenol (1 volume of phenol: glycerol: lactic acid: water (1:1:1:1) and 2 volumes of ethanol). Leaves were transferred to fresh alcoholic lactophenol and incubated overnight at room temperature. After two washes with 50% ethanol, leaves were stained with aniline blue and callose deposits were visualized under a fluorescence microscope. Quantification of callose deposits was performed using the Fiji distribution of ImageJ.

#### Confocal microscopy

Two-week old *Arabidopsis* GFP-HDEL plants were vacuum infiltrated with either water or 100 nM flg22. Three hours after the treatment, cotyledons were detached and observed under a Zeiss LSM710 confocal microscope equipped with a LDC-apochromat 40x/1.1W Korr M27 water-immersion objective (NA 1.1). GFP was excited at 488 nm and emission collected at 500-550 nm. For monitoring of the dynamics of the ER bodies, cotyledons were treated as above and visualized using a Leica SP8 confocal microscope (Leica Microsystems GmbH, Wetzlar, Germany). GFP was excited at 488 nm and emission collected at 500-550 nm.

For visualization of bacterial-induced ER bodies, five-week old *Arabidopsis* GFP-HDEL plants were brush-inoculated with a 5×10^7^ CFU/mL suspension of the indicated strain (Fig. 3A). Three days after inoculation, leaves were observed as described above.

For visualization of bacterial microcolonies, five-week old *Arabidopsis* GFP-HDEL plants were syringe infiltrated with a 5×10^4^ CFU/mL suspension and leaves were visualized using the Leica SP5 II confocal microscope (Leica Microsystems GmbH, Wetzlar, Germany). Images of eYFP and GFP were sequentially obtained using the following conditions (excitation/ emission): eYFP (514 nm/ 525 to 600 nm), GFP (488/ 500 to 525 nm). Z series imaging was taken at 1 μm interval. Image processing was performed using Leica LAS AF (Leica Microsystems). ER bodies were quantified using Fiji distribution of ImageJ software.

#### *S. exigua* performance

*S. exigua* eggs were kindly provided by Dr. Salvador Herrero (U. Valencia). After eclosion, *S. exigua* larvae were fed with an artificial diet (Elvira et al., 2010). Three days after eclosion, five larvae of approximately same size were transferred to a four-to-five week old *Arabidopsis* plant grown in Jiffy-7 pods. Larvae were transfer to new plants every 24 hours to avoid food privation. Seven days after plant feeding, fresh weight of the larvae was measured. For each experiment, five replicates were performed (25 larvae per genotype).

## ACKNOWLEDGMENTS

J.S. Rufián has been supported by a FPI fellowship associated with a grant of E.R. Bejarano (MICINN, Spain; AGL2010-22287-C02-2), funds from BIO2012-35641 (MICINN, Spain) granted to C.R. Beuzón, and Plan Propio de la Universidad de Málaga – Andalucía Tech. The work was co-funded by European Regional Development Funds (FEDER). G. Coaker is supported by grants from the National Institutes of Health (RO1GM092772, R35GM13640). We thank Alberto Macho for hosting J.S. Rufián in his research laboratory and providing feedback on the manuscript.

## Supplemental Figure Legends

**Figure S1. Downregulation of *NAI1* and *PYK10* after flg22 treatment**.

Expression pattern of *NAI1* and *PYK10* after flg22 treatment (5, 10, 30, 90 and 180 minutes) in Col-0 and *fls2* plants. Data were obtained from Bjornson et al. (2021). Fourteen-day old Arabidopsis Col-0 or *fls2* seedlings were immersed in 1 μM flg22 and samples were taken at the indicated time points.

**Figure S2. *bglu18-1* and *pbp1-1* show enhanced resistance to *P. syringae***

(**A**) Flg22-induced ROS burst in Col-0, *bglu18-1*, and *pbp1-1*. Three-week-old *Arabidopsis* leaf discs were treated with 100 nM flg22 and ROS was quantified using a luminescence based assay. Error bars indicate ±SE, *n* = 16. The experiment was repeated twice with similar results and a representative experiment is shown. (**B**) Growth of *P. syringae* pv. *tomato* DC3000 in Col-0, *bglu18-1*, and *pbp1-1*. Five-week-old plant leaves were syringe inoculated with a suspension of 5×10^4^ CFU/mL. Four days after inoculation, bacteria were recovered and quantified. Values for each individual plant are shown (*n*=6). Bars represent the mean value ± SE. Statistically differences were determined with one-way ANOVA (α= 0.05) with Tukey’s multiple comparisons test and different letters indicate statistical significance. The experiment was repeated twice with similar results and a representative experiment is shown.

**Figure S3. Coronatine induce the expression of ER body and indole glucosinolate related genes**.

(A) Transcriptomic analysis of several genes related with ER bodies formation or indole-glucosinolate metabolism. Bars show Log_2_ signal values from microarrays datasets. Data was obtained from the study by Thilmony et al. (2006), available in Genevestigator. *Arabidopsis* Col-5 leaves were inoculated with water (Mock), or a 10^6^ cfu/ml suspension of *Pst* DC3000 or *Pst* DC3118 (cor^-^). Samples were taken 24 hours after inoculation. (B) Coexpression analyses of PYK10 (At3G09260). Analyses were performed using ATTED-II software.

## Literature Cited

Bednarek, P., Piślewska-Bednarek, M., Svatos, A., Schneider, B., Doubsky, J., Mansurova, M., Humphry, M., Consonni, C., Panstruga, R., Sanchez-Vallet, A., Molina, A., and Schulze-Lefert, P. (2009). A glucosinolate metabolism pathway in living plant cells mediates broad-spectrum antifungal defense. In Science (American Association for the Advancement of Science), pp. 101–106.

Bender, C.L., Alarcón-Chaidez, F., and Gross, D.C. 1999. Pseudomonas syringae phytotoxins: mode of action, regulation, and biosynthesis by peptide and polyketide synthetases. Microbiol Mol Biol Rev 63:266–292.

Bertani, G. 1951. Studies on lysogenesis. I. The mode of phage liberation by lysogenic Escherichia coli. Journal of bacteriology 62:293–300.

Bjornson, M., Pimprikar, P., Nurnberger, T., and Zipfel, C. 2021. The transcriptional landscape of Arabidopsis thaliana pattern-triggered immunity. Nat Plants.

Brooks, D.M., Hernandez-Guzman, G., Kloek, A.P., Alarcon-Chaidez, F., Sreedharan, A., Rangaswamy, V., Penaloza-Vazquez, A., Bender, C.L., and Kunkel, B.N. 2004. Identification and characterization of a well-defined series of coronatine biosynthetic mutants of Pseudomonas syringae pv. tomato DC3000. Mol Plant Microbe Interact 17:162–174.

Chung, H.S., Koo, A.J.K., Gao, X., Jayanty, S., Thines, B., Jones, A.D., and Howe, G.A. (2008). Regulation and function of Arabidopsis JASMONATE ZIM-domain genes in response to wounding and herbivory. In Plant physiology (American Society of Plant Biologists), pp. 952–964.

Cipollini, D., Enright, S., Traw, M.B., and Bergelson, J. (2004). Salicylic acid inhibits jasmonic acid-induced resistance of Arabidopsis thaliana to Spodoptera exigua. In Molecular Ecology, pp. 1643–1653.

Clay, N.K., Adio, A.M., Denoux, C., Jander, G., and Ausubel, F.M. 2009. Glucosinolate metabolites required for an Arabidopsis innate immune response. Science 323:95–101.

Couto, D., and Zipfel, C. 2016. Regulation of pattern recognition receptor signalling in plants. Nature reviews. Immunology 16:537–552.

Cuppels, D.A. 1986. Generation and Characterization of Tn5 Insertion Mutations in Pseudomonas syringae pv. tomato. Appl. Environ. Microbiol. 51:323–327.

Elvira, S., Gorria, N., Munoz, D., Williams, T., and Caballero, P. 2010. A simplified low-cost diet for rearing Spodoptera exigua (Lepidoptera: Noctuidae) and its effect on S. exigua nucleopolyhedrovirus production. J Econ Entomol 103:17–24.

Fuchs, R., Kopischke, M., Klapprodt, C., Hause, G., Meyer, A.J., Schwarzländer, M., Fricker, M.D., and Lipka, V. (2016). Immobilized Subpopulations of Leaf Epidermal Mitochondria Mediate PENETRATION2-Dependent Pathogen Entry Control in Arabidopsis. In The Plant Cell … (American Society of Plant Biologists), pp. 130–145.

Geng, X., Cheng, J., Gangadharan, A., and Mackey, D. (2012). The coronatine toxin of Pseudomonas syringae is a multifunctional suppressor of Arabidopsis defense. In The Plant Cell …, pp. 4763–4774.

Ghorbel, M., Brini, F., Sharma, A., and Landi, M. 2021. Role of jasmonic acid in plants: the molecular point of view. Plant Cell Rep.

Gimenez-Ibanez, S., Boter, M., Ortigosa, A., García-Casado, G., Chini, A., Lewsey, M.G., Ecker, J.R., Ntoukakis, V., and Solano, R. (2017). JAZ2 controls stomata dynamics during bacterial invasion. In New Phytol., pp. 1378–1392.

Gómez-Gómez, L., and Boller, T. 2000. FLS2: an LRR receptor-like kinase involved in the perception of the bacterial elicitor flagellin in Arabidopsis. Mol Cell 5:1003–1011.

Guo, Q., Yoshida, Y., Major, I.T., Wang, K., Sugimoto, K., Kapali, G., Havko, N.E., Benning, C., and Howe, G.A. 2018. JAZ repressors of metabolic defense promote growth and reproductive fitness in Arabidopsis. Proc Natl Acad Sci U S A 115:E10768–E10777.

Henry, E., Yadeta, K.A., and Coaker, G. 2013. Recognition of bacterial plant pathogens: local, systemic and transgenerational immunity. New Phytol 199:908–915.

Johansson, O.N., Fantozzi, E., Fahlberg, P., Nilsson, A.K., Buhot, N., Tör, M., and Andersson, M.X. (2014). Role of the penetration-resistance genes PEN1, PEN2and PEN3in the hypersensitive response and race-specific resistance in Arabidopsis thaliana. In Plant J., pp. 466–476.

Lahrmann, U., Strehmel, N., Langen, G., Frerigmann, H., Leson, L., Ding, Y., Scheel, D., Herklotz, S., Hilbert, M., and Zuccaro, A. 2015. Mutualistic root endophytism is not associated with the reduction of saprotrophic traits and requires a noncompromised plant innate immunity. New Phytol 207:841–857.

Lipka, V., Dittgen, J., Bednarek, P., Bhat, R., Wiermer, M., Stein, M., Landtag, J., Brandt, W., Rosahl, S., Scheel, D., Llorente, F., Molina, A., Parker, J., Somerville, S., and Schulze-Lefert, P. (2005). Pre-and postinvasion defenses both contribute to nonhost resistance in Arabidopsis. In Science (American Association for the Advancement of Science), pp. 1180–1183.

Macho, A.P. 2015. Subversion of plant cellular functions by bacterial type-III effectors: beyond suppression of immunity. New Phytol.

Matsushima, R., Fukao, Y., Nishimura, M., and Hara-Nishimura, I. (2004). NAI1 gene encodes a basic-helix-loop-helix-type putative transcription factor that regulates the formation of an endoplasmic reticulum-derived structure, the ER body. In THE PLANT CELL ONLINE (American Society of Plant Biologists), pp. 1536–1549.

Matsushima, R., Hayashi, Y., Kondo, M., Shimada, T., Nishimura, M., and Hara-Nishimura, I. (2002). An endoplasmic reticulum-derived structure that is induced under stress conditions in Arabidopsis. In Plant physiology (American Society of Plant Biologists), pp. 1807–1814.

Matsushima, R., Hayashi, Y., Yamada, K., Shimada, T., Nishimura, M., and Hara-Nishimura, I. (2003). The ER body, a novel endoplasmic reticulum-derived structure in Arabidopsis. In Plant Cell Physiol., pp. 661–666.

Melotto, M., Underwood, W., and He, S.Y. 2008. Role of stomata in plant innate immunity and foliar bacterial diseases. Annu Rev Phytopathol 46:101–122.

Melotto, M., Underwood, W., Koczan, J., Nomura, K., and He, S.Y. 2006. Plant stomata function in innate immunity against bacterial invasion. Cell 126:969–980.

Millet, Y.A., Danna, C.H., Clay, N.K., and Songnuan, W. (2010). Innate immune responses activated in Arabidopsis roots by microbe-associated molecular patterns. In The Plant Cell ….

Mitsuhashi, N., Shimada, T., Mano, S., Nishimura, M., and Hara-Nishimura, I. 2000. Characterization of organelles in the vacuolar-sorting pathway by visualization with GFP in tobacco BY-2 cells. Plant Cell Physiol 41:993–1001.

Müller, R., de Vos, M., Sun, J.Y., Sønderby, I.E., Halkier, B.A., Wittstock, U., and Jander, G. (2010). Differential Effects of Indole and Aliphatic Glucosinolates on Lepidopteran Herbivores. In J Chem Ecol, pp. 905–913.

Nagano, A.J., Matsushima, R., and Hara-Nishimura, I. (2005). Activation of an ER-body-localized beta-glucosidase via a cytosolic binding partner in damaged tissues of Arabidopsis thaliana. In Plant Cell Physiol., pp. 1140–1148.

Nagano, A.J., Fukao, Y., Fujiwara, M., Nishimura, M., and Hara-Nishimura, I. (2008). Antagonistic jacalin-related lectins regulate the size of ER body-type beta-glucosidase complexes in Arabidopsis thaliana. In Plant Cell Physiol., pp. 969–980.

Nagano, A.J., Maekawa, A., Nakano, R.T., Miyahara, M., Higaki, T., Kutsuna, N., Hasezawa, S., and Hara-Nishimura, I. (2009). Quantitative analysis of ER body morphology in an Arabidopsis mutant. In Plant Cell Physiol., pp. 2015–2022.

Nakano, R.T., Yamada, K., Bednarek, P., Nishimura, M., and Hara-Nishimura, I. 2014a. ER bodies in plants of the Brassicales order: biogenesis and association with innate immunity. Front Plant Sci 5:73.

Nakano, R.T., Yamada, K., Bednarek, P., Nishimura, M., and Hara-Nishimura, I. (2014b). ER bodies in plants of the Brassicales order: biogenesis and association with innate immunity. In Front. Plant Sci. (Frontiers), pp. 73.

Nakano, R.T., Piślewska-Bednarek, M., Yamada, K., Edger, P.P., Miyahara, M., Kondo, M., Böttcher, C., Mori, M., Nishimura, M., Schulze-Lefert, P., Hara-Nishimura, I., and Bednarek, P. (2017). PYK10 myrosinase reveals a functional coordination between endoplasmic reticulum bodies and glucosinolates in Arabidopsis thaliana. In Plant J., pp. 204–220.

Nakazaki, A., Yamada, K., Kunieda, T., Tamura, K., Hara-Nishimura, I., and Shimada, T. (2019a). Biogenesis of leaf endoplasmic reticulum body is regulated by both jasmonate-dependent and independent pathways. In Plant Signaling & Behavior (Taylor & Francis), pp. 1–3.

Nakazaki, A., Yamada, K., Kunieda, T., Sugiyama, R., Hirai, M.Y., Tamura, K., Hara-Nishimura, I., and Shimada, T. (2019b). Leaf Endoplasmic Reticulum Bodies Identified in Arabidopsis Rosette Leaves Are Involved in Defense against Herbivory. In Plant physiology, pp. 1515–1524.

Ogasawara, K., Yamada, K., Christeller, J.T., Kondo, M., Hatsugai, N., Hara-Nishimura, I., and Nishimura, M. (2009). Constitutive and inducible ER bodies of Arabidopsis thaliana accumulate distinct beta-glucosidases. In Plant Cell Physiol., pp. 480–488.

Rehrig, E.M., Appel, H.M., Jones, A.D., and Schultz, J.C. (2014). Roles for jasmonate-and ethylene-induced transcription factors in the ability of Arabidopsis to respond differentially to damage caused by two insect herbivores. In Front. Plant Sci. (Frontiers), pp. 407.

Rufian, J.S., Macho, A.P., Corry, D.S., Mansfield, J.W., Ruiz-Albert, J., Arnold, D.L., and Beuzon, C.R. 2018. Confocal microscopy reveals in planta dynamic interactions between pathogenic, avirulent and non-pathogenic Pseudomonas syringae strains. Molecular plant pathology 19:537–551.

Santamaría, M.E., Martinez, M., Arnaiz, A., Ortego, F., Grbic, V., and Diaz, I. (2017). MATI, a Novel Protein Involved in the Regulation of Herbivore-Associated Signaling Pathways. In Front. Plant Sci., pp. 80–18.

Sherameti, I., Venus, Y., Drzewiecki, C., Tripathi, S., Dan, V.M., Nitz, I., Varma, A., Grundler, F.M., and Oelmüller, R. (2008). PYK10, a beta-glucosidase located in the endoplasmatic reticulum, is crucial for the beneficial interaction between Arabidopsis thaliana and the endophytic fungus Piriformospora indica. In Plant J. (Blackwell Publishing Ltd), pp. 428–439.

Sugiyama, R., and Hirai, M.Y. (2019). Atypical Myrosinase as a Mediator of Glucosinolate Functions in Plants. In Front. Plant Sci. (Frontiers), pp. 1008.

Thilmony, R., Underwood, W., and He, S.Y. 2006. Genome-wide transcriptional analysis of the Arabidopsis thaliana interaction with the plant pathogen Pseudomonas syringae pv. tomato DC3000 and the human pathogen Escherichia coli O157:H7. Plant J 46:34–53.

Van Oosten, V.R., Bodenhausen, N., Reymond, P., Van Pelt, J.A., Van Loon, L.C., Dicke, M., and Pieterse, C.M.J. (2008). Differential effectiveness of microbially induced resistance against herbivorous insects in Arabidopsis. In Mol. Plant Microbe Interact. (The American Phytopathological Society), pp. 919–930.

Weiler, E.W., Kutchan, T.M., Gorba, T., Brodschelm, W., Niesel, U., and Bublitz, F. 1994. The Pseudomonas phytotoxin coronatine mimics octadecanoid signalling molecules of higher plants. FEBS letters 345:9–13.

Wittstock, U., and Burow, M. (2010). Glucosinolate Breakdown in Arabidopsis: Mechanism, Regulation and Biological Significance. In The Arabidopsis Book, pp. e0134–0114.

Wittstock, U., Kurzbach, E., Herfurth, A.M., and Stauber, E.J. (2016). Glucosinolate Breakdown. In Advances in Botanical Research (Elsevier Ltd), pp. 125–169.

Xu, Z., Escamilla-Trevino, L., Zeng, L., Lalgondar, M., Bevan, D., Winkel, B., Mohamed, A., Cheng, C.L., Shih, M.C., Poulton, J., and Esen, A. 2004. Functional genomic analysis of Arabidopsis thaliana glycoside hydrolase family 1. Plant Mol Biol 55:343–367.

Yamada, K., Hara-Nishimura, I., and Nishimura, M. (2011). Unique defense strategy by the endoplasmic reticulum body in plants. In Plant Cell Physiol., pp. 2039–2049.

Yamada, K., Nagano, A.J., Nishina, M., Hara-Nishimura, I., and Nishimura, M. (2008). NAI2 is an endoplasmic reticulum body component that enables ER body formation in Arabidopsis thaliana. In THE PLANT CELL ONLINE (American Society of Plant Biologists), pp. 2529–2540.

