## Supplemental Fig 1 for "ER bodies are induced by *Pseudomonas syringae* and negatively regulate immunity"

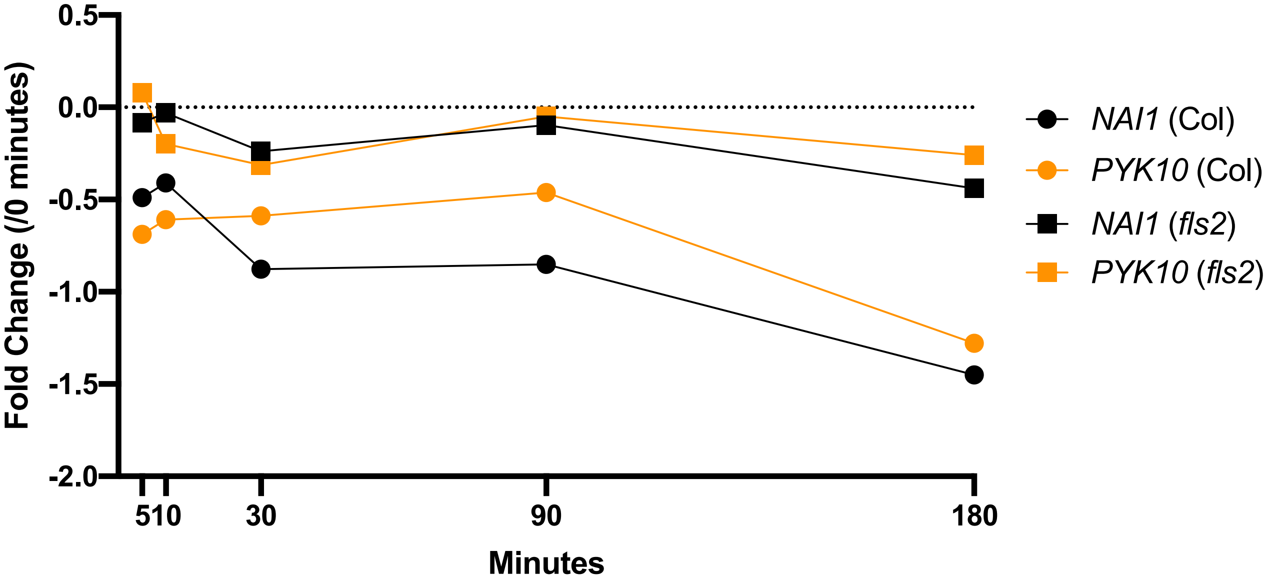


**Figure S1. Downregulation of *NAI1* and *PYK10* after flg22 treatment**
Expression pattern of *NAI1* and *PYK10* after flg22 treatment (5, 10, 30, 90 and 180 minutes) in Col-0 and *fls2* plants. Data were obtained from Bjornson et al. (2021). Fourteen-day old Arabidopsis Col-0 or *fls2* seedlings were inmersed in 1 μM flg22 and samples were taken at the indicated time points.
