## Supplemental Fig 2 for "ER bodies are induced by *Pseudomonas syringae* and negatively regulate immunity"

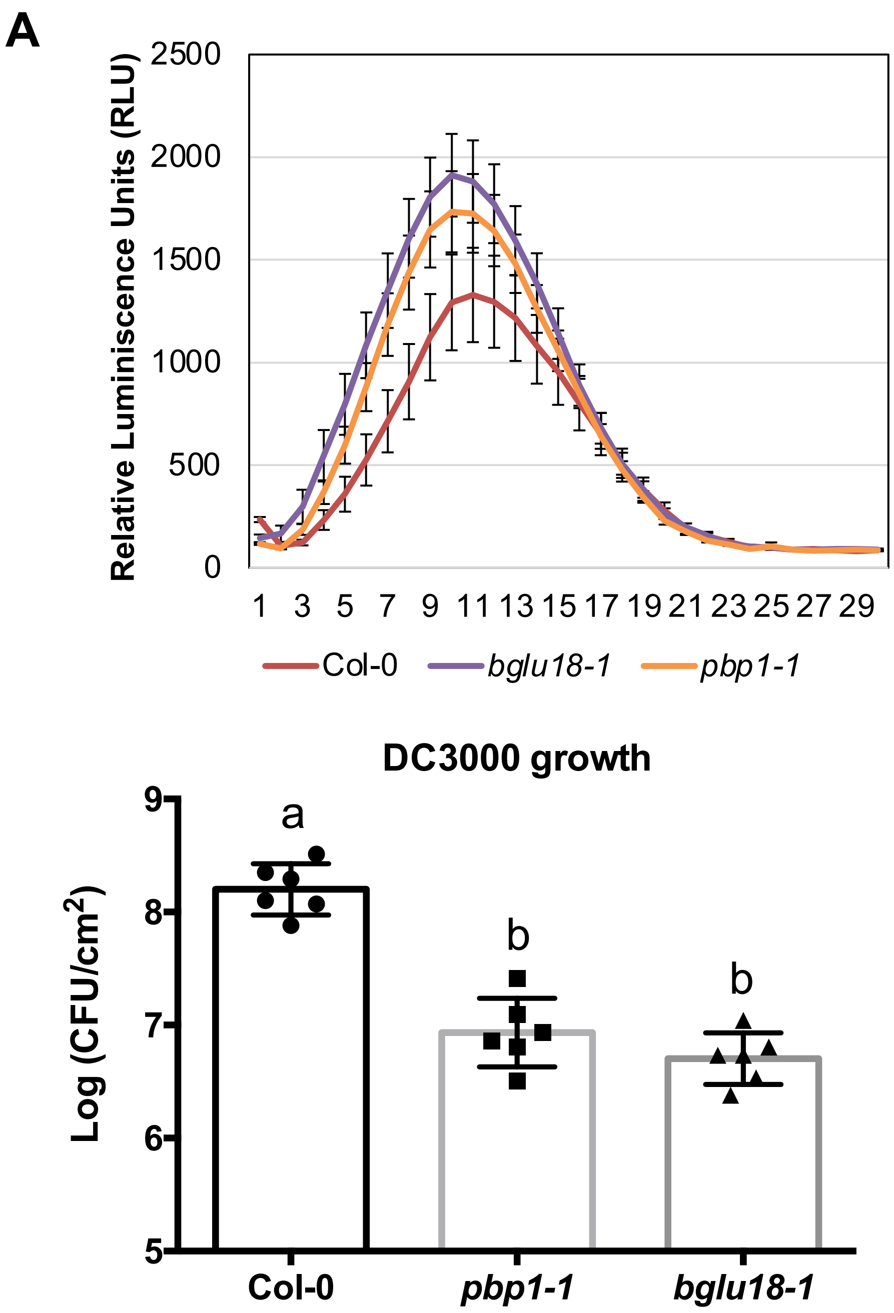


**Figure S2. *bglu18-1* and *pbp1-1* also show enhanced resistance to *P. syringae***

(**A**) flg22-induced ROS burst in Col-0, *bglu18-1*, and *pbp1-1*. Three-week-old Arabidopsis leaf discs were treated with 100 nM flg22 and ROS was quantified using a luminescence based assay. Error bars indicate SE. *n* = 16. The experiment was repeated twice with similar results and a representative experiment is shown. (**B**) Growth of *P. syringae* pv. tomato DC3000 in Col-0, *bglu18-1*, and *pbp1-1*. Five-week-old plant leaves were syringe inoculated with a suspension of $5x10^4$ CFU/mL. Four days after inoculation, bacteria were recovered and quantified. Values for each individual plant are shown (*n*=6). Bars represent the mean value ± SE. Statistically differences were determined with one-way ANOVA (α= 0.05) with Tukey's multiple comparisons test and different letters indicate statistical significance. The experiment was repeated twice with similar results and a representative experiment is shown.
