## Supplemental Fig 3 for "ER bodies are induced by *Pseudomonas syringae* and negatively regulate immunity"

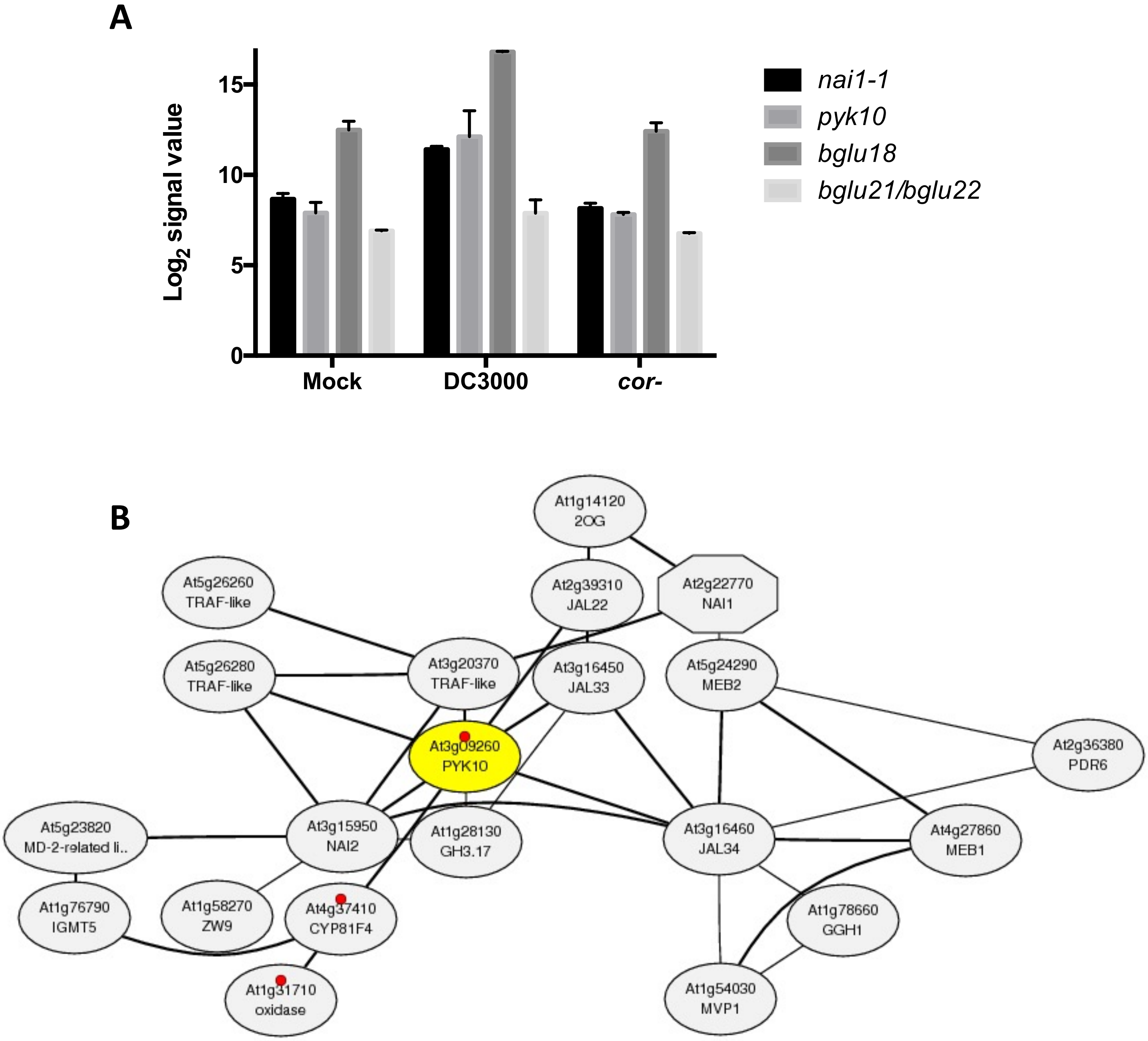


**Figure S3. Coronatine induce the expression of ER bodies- and indole glucosinolates-related genes.** (A) Transcriptomic analysis of several genes related with the ER bodies or the indole-glucosinolates metabolism. Bars show Log2 signal values from microarrays datasets. Data was obtained from the study by Thilmony et al. (2006), available in Genevestigator. Arabidopsis Col-5 leaves were inoculated with water (Mock), or a 106 cfu/ml suspension of DC3000 or DC3118 (cor-). Samples were taken 24 hours after inoculation. (B) Coexpression analysis of PYK10 (At3G09260). Analysis was made using ATTED-II software.
